# LineageFlow: growth-corrected state dynamics for probabilistic cell-fate prediction

**DOI:** 10.64898/2026.10.17.759846

**Authors:** Yang Liu, Qi Liu, Wei Wang, Jian Liu, Yu Wang, Wei Zhang

## Abstract

Single-cell lineage tracing links early transcriptional states to the fates of their descendants, but forecasting clonal fate remains challenging because differentiation and unequal cell expansion jointly shape descendant populations. Here we introduce LineageFlow, a computational framework that combines continuous-time cell-state transitions, relative net growth and state-conditioned generative flows to predict future fate distributions and transcriptional profiles from early single-cell measurements. In clone-disjoint evaluations of lineage-traced hematopoiesis, LineageFlow improves fate prediction and expression-distribution accuracy over matched computational baselines. A separately retrained analysis of cord-blood lineage data shows a similar advantage at a longer prediction horizon. Controlled simulations show that accounting for differential growth improves fate prediction when descendant states expand at different rates, while offering little benefit under equal growth. These findings highlight the importance of modeling state transitions and relative expansion together when predicting the fate of unseen clones.

## 1 Introduction

Understanding how an early cell population gives rise to distinct differentiated progeny is a central problem in developmental and stem-cell biology. Single-cell RNA sequencing reveals molecular heterogeneity, while lineage tracing connects cells that share ancestry across experimental time points [1–3]. CoSpar estimates early fate bias, LineageOT incorporates ancestry into temporal transport, and CLADES and DestinyNet study clonal dynamics and fate prediction [4–7]. CellRank 2 offers a complementary probabilistic fate-mapping framework [8]. Genetic barcoding and scar-based recording have made it possible to read out both cell identity and ancestry across diverse developmental systems [9–14]. Together, these measurements offer a way to investigate how early transcriptional states relate to later cell fates. The prediction problem, however, is not the same as reconstructing a continuous trajectory for an individual cell. Sequencing is destructive, barcodes identify sampled relatives, and the eventual descendants of the measured progenitor are not observed directly. A useful prospective forecast must therefore describe the possible states of sampled descendants, together with uncertainty, at a specified future time.

An important difficulty is that descendant proportions reflect both *differentiation* and *population expansion*. A lineage may become enriched in a particular state because cells enter that state more frequently, because cells already in that state divide faster or survive longer, or because of both mechanisms. These explanations need not be distinguishable from normalized state fractions alone, a confounding problem also recognized in models of population flux from single-cell time courses [15, 16]. In particular, a continuous-time Markov chain (CTMC) describes movement of conserved probability mass between states; it does not, without modification, account for differences in the number of descendants contributed by those states. This distinction becomes especially relevant as forecasting horizons increase. Growth-aware single-cell modeling provides an additional point of comparison through PRESCIENT and TIGON [17, 18]. For the distinct task of generating within-fate transcriptional profiles, conditional flow matching supplies a generative formulation [19]; CTMC–flow coupling was previously explored in a nonbiological setting in Sketch2Colab [20].

Several approaches have transformed the analysis of singlecell dynamics, but they address different observation and inference settings. Trajectory-inference and fate-mapping methods organize cells along developmental paths or estimate terminal-state propensities [8, 21–26]. RNA velocity and vector-field approaches infer local expression dynamics rather than directly observing expansion of barcode-defined descendants [27–30]. CoSpar combines transcriptional and clonal information to recover fate bias from sparse lineage observations [4]; the more recent DestinyNet addresses fate clustering, flow and prediction from lineage-traced single-cell profiles [7]. Temporal transport and lineage-aware optimal-transport approaches link measurements across time points [5, 31–33]. Population-based methods incorporate growth, differentiation kinetics or clone-specific behaviour [6, 17, 18, 34]. These methods provide essential points of comparison, but many are designed to analyze a complete observed dataset rather than predict the future of an unseen clone from its early measurements alone. Their native tasks are therefore not all directly interchangeable.

Here we introduce LineageFlow, a model for *finite-horizon, clone-disjoint prediction* of descendant fate and transcriptional state. The framework has two coupled components: a clone-conditioned state model that propagates expected lineage mass under state transitions and relative net growth, and a state-conditioned flow model that generates continuous expression profiles. We deliberately maintain a distinction between biological time, which indexes the descendant forecast, and the interpolation time used to generate expression. Our adaptation of that CTMC–flow architecture changes the state and observation space, adds relative-growth propagation, and evaluates the resulting predictions against lineage-resolved biological measurements. It does not claim that the CTMC–flow pairing is itself new.

We assess the approach using early-to-late predictions in the LARRY hematopoiesis time course, with barcode-connected components held out in their entirety [1]. We compare fate probabilities and generated expression against direct conditional and time-dependent alternatives and against training-only prospective adaptations of published approaches. Additional analyses examine rare cell states, uneven descendant sampling, model components, transition-graph assumptions and a separately retrained cord-blood lineage dataset. Finally, controlled five-state systems isolate settings in which differential growth should matter. Together, these analyses ask a focused question: when does explicitly representing statedependent descendant expansion improve the prediction of future clonal composition?

## 2 Model framework

### 2.1 A growth-aware framework predicts both descendant composition and expression

LineageFlow takes an early clone as input and returns a probability distribution over the states and transcriptomic profiles of its descendants at a chosen future horizon (Fig. 1). Early observations are summarized by their transcriptional features, their annotated state counts and the experimental starting population. Barcode identities establish which cells belong to the same clone-connected component but are not supplied as predictive features. The model thus learns from many training clones without relying on a test clone’s barcode as an identifier.

**Figure 1.**
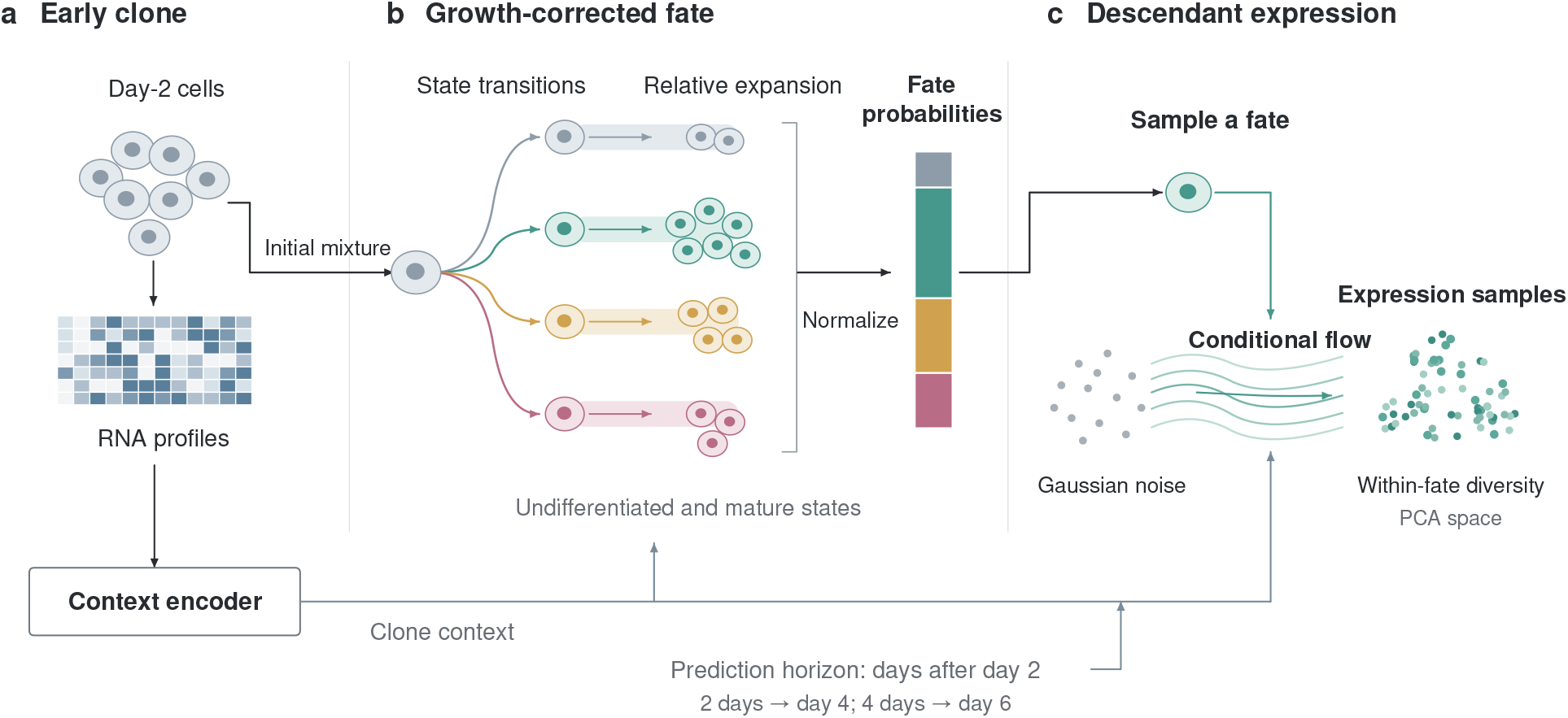
Overview of clone-conditional descendant prediction. **a**, Day-2 cells provide early transcriptional profiles, initial state proportions and starting-population information. **b**, A context-dependent state-transition generator and relative net-growth vector evolve expected lineage mass; normalization yields the predicted descendant-state mixture. Three mature states are drawn for clarity, although the primary analysis uses ten mature states and an undifferentiated state. **c**, A state is sampled from that mixture before a conditional continuous flow generates an expression profile in 50-dimensional principal-component space. Biological horizons of 2 and 4 days correspond to observations on days 4 and 6. Only training clones provide future-state and expression targets; test-clone forecasts use early measurements alone. The CTMC-guided flow organization is adapted from Sketch2Colab [20].

The first module learns which state transitions are permitted and how their rates depend on the early clone. A second set of parameters describes relative net expansion in each state. Rather than normalizing state-transition probabilities directly, the model evolves nonnegative expected lineage mass and then converts that mass to descendant proportions. This produces a forecast that accounts for both the appearance of new states and the unequal abundance of their progeny. The second module generates a transcriptomic profile conditional on a fate sampled from that forecast. This ordering preserves discrete alternative fates while allowing continuous expression variability within each fate; it does not average expression trajectories across incompatible terminal states.

The population component is expressed compactly as

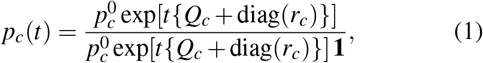

where 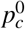 is the smoothed early-state distribution of clone *c, Q*_*c*_ is a valid CTMC transition generator and *r*_*c*_ contains relative net-growth terms. When all states grow at the same rate, the common growth factor cancels and Eq. 1 reduces to ordinary CTMC occupancy. With unequal growth, the two models generally predict different descendant compositions. Crucially, the quantity in Eq. 1 is *normalized expected mass*, not an exact likelihood for stochastic branching counts or for the capture process. Model fitting and the state-conditional flow are described in Methods.

### 2.2 Prediction target and clone representation

We consider discrete observations of cell states and gene expression from multiple experimentally sampled time points. The unit of prediction is a barcode-connected clone component, indexed by *c*. Let 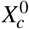 denote expression profiles from its early sampled cells, 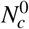the early sample size and 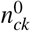 the early count assigned to annotated state ∈ {*k* 1, …, *K*}. For a future biological horizon *t*, the labelled descendant observations have state counts *y*_*c*_(*t*) and, if *N*_*c*_(*t*) = ∑_*k*_ *y*_*ck*_(*t*) *>* 0, empirical proportions *f*_*c*_(*t*) = *y*_*c*_(*t*)*/N*_*c*_(*t*). The target is the state of a randomly selected labelled descendant at that horizon, together with its conditional distribution of transcriptional profiles.

Initial state proportions are estimated with additive smoothing,

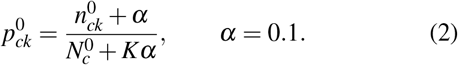

The smoothing prevents exact zeros when an annotated state is absent from a small early sample, at the cost of assigning some initial mass to unobserved states. Consequently, we explicitly examine pseudocount sensitivity (Fig. 3a). The expression representation is constructed from 2,000 highly variable genes and 50 principal components; all gene selection, preprocessing and PCA fitting use training cells only. We summarize each early component by the mean and variance of its PCA coordinates and its starting-population category. A shared neural encoder maps these summaries to clone context *h*_*c*_. Barcode strings are used to establish group membership and splits, never as predictive features. In the LARRY analysis, *t* = 2 and 4 days corresponds to outcomes measured on experimental days 4 and 6, with day 2 as the prediction origin.

**Figure 2.**
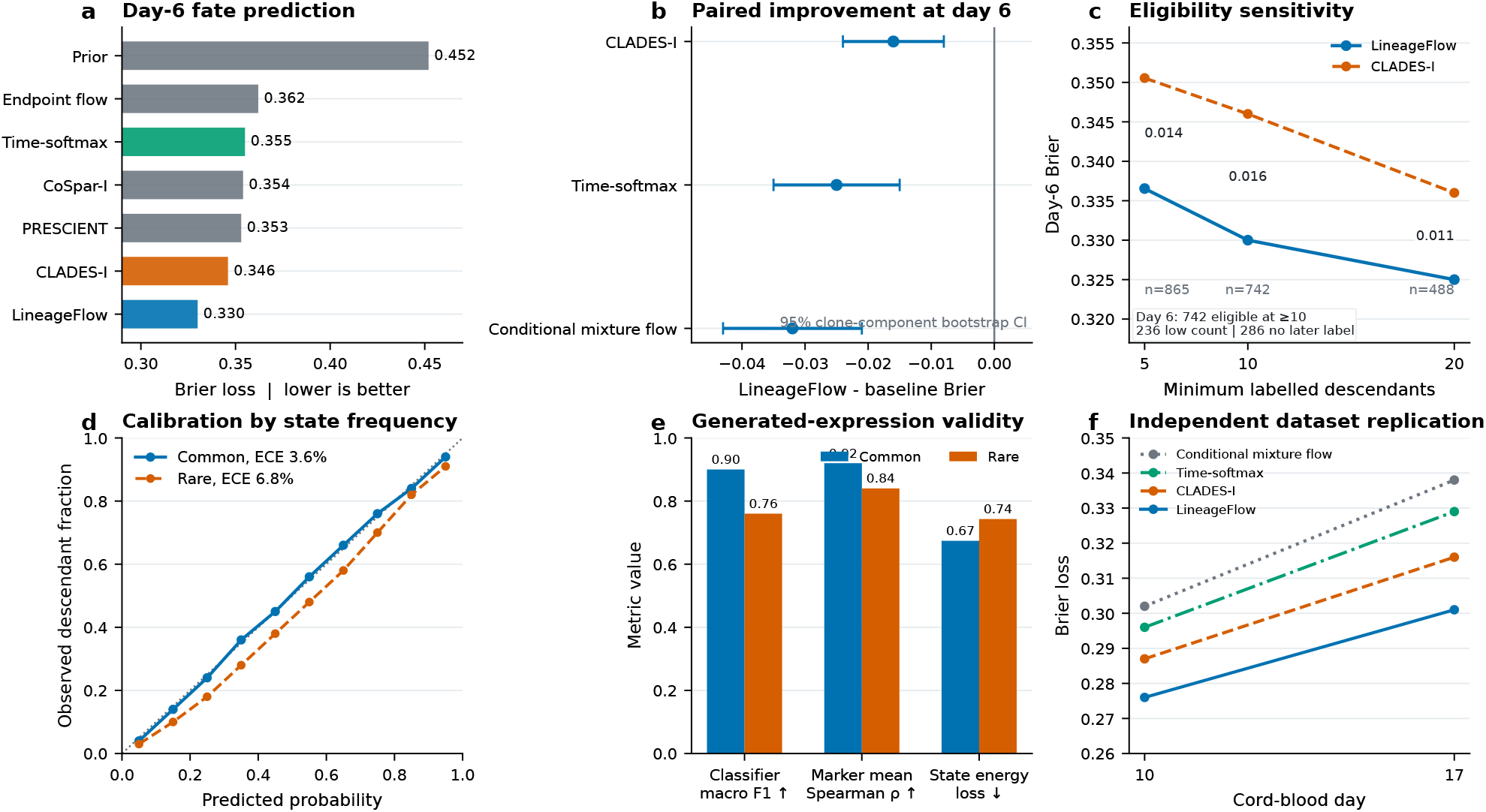
Predictive performance, cohort sensitivity and expression diagnostics in lineage-traced data. **a**, Day-6 categorical Brier loss for representative models, with lower values indicating better predicted descendant-state probabilities. **b**, Paired LineageFlow-minus-comparator Brier differences and 95% bootstrap intervals from held-out clone components; negative differences favor LineageFlow. **c**, Sensitivity to the minimum observed descendant count, with day-6 cohort accounting in the inset. **d**, Reliability curves separately summarize common and rare states. **e**, Generated-expression diagnostics compare classifier agreement, normalized marker means and state-conditional energy loss. **f**, Day-10 and day-17 prediction after retraining all compared models on an independent cord-blood lineage dataset. All biological evaluations use future observations for scoring, not for conditioning held-out predictions.

**Figure 3.**
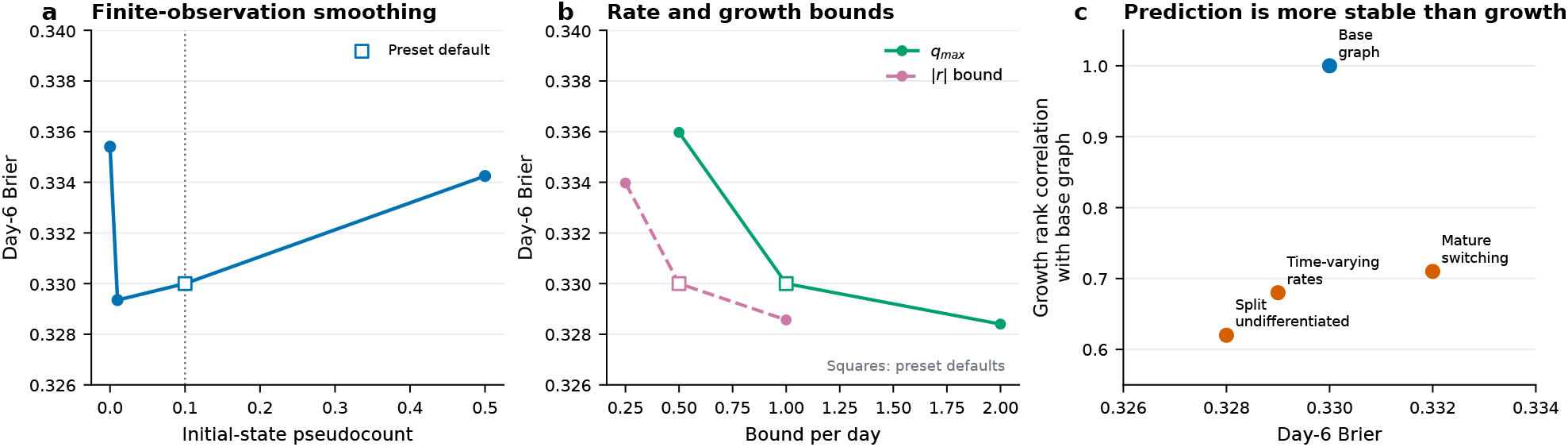
Sensitivity of outcome predictions to modeling assumptions. **a**, Day-6 Brier loss as the initial-state pseudocount is varied. **b**, Rate and growth-bound sensitivity sweeps; marked defaults are retained even where another value yields a lower test score. **c**, Allowing mature-state switching, splitting the undifferentiated state or using time-varying rates alters fitted relative-growth rankings more strongly than predictive Brier loss. Each sweep varies a single assumption while keeping the component partitions and training recipe fixed.

### 2.3 Valid state transitions and relative net growth

We represent discrete changes in annotated fate state by a clone-conditioned, time-homogeneous CTMC generator *Q*_*c*_. A masked neural head assigns each permitted transition from state *i* to state *j* the rate

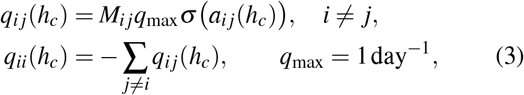

where *M*_*ij*_ ∈ {0, 1} encodes permitted edges and *σ* is the logistic sigmoid. Off-diagonal entries are nonnegative and row sums vanish, so *Q*_*c*_ is a valid CTMC generator. Related continuous-time and discrete-flow constructions have been used in generative modeling [35, 36]; here the state generator instead evolves biological-time descendant mass. In the primary LARRY graph the undifferentiated category may enter any of ten mature states; the mature categories are absorbing *with respect to state transitions*. This does not prohibit their number of descendants from changing. Alternative graph structures are evaluated separately.

A second neural head produces a clone-specific vector *r*_*c*_ = (*r*_*c*1_, …, *r*_*cK*_) of bounded relative net-growth terms. The default bound is |*r*_*ck*_| ≤ 0.5 day^−1^, and the vector is zerocentered so **1**^⊤^*r*_*c*_ = 0. A common shift in all growth rates cannot be identified from normalized composition because it cancels on normalization. Accordingly, *r*_*c*_ is interpreted as *relative effective expansion*, not as an independent estimate of absolute proliferation, survival or death.

For the initial probability row vector 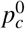, we evolve expected mass under the linear operator

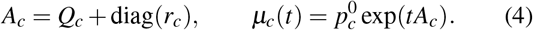

The predicted descendant-state composition is *p*_*c*_(*t*) = *µ*_*c*_(*t*)*/*[*µ*_*c*_(*t*)**1**] (Eq. 1). The matrix *A*_*c*_ has nonnegative offdiagonal entries; hence its exponential preserves nonnegative mass for nonnegative initial conditions. Differentiation of the normalized expression gives

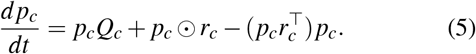

Here⊙ denotes elementwise multiplication. The first term changes the state mixture through transitions; the final two terms change relative composition through differences in net expansion. If *r*_*c*1_ =···= *r*_*cK*_, those growth-dependent terms cancel and the forecast reduces to 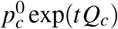. Equation 5 also clarifies why rate identification is difficult: different combinations of transition structure and relative expansion may induce similar changes in normalized fractions.

### 2.4 Conditional flow for transcriptomic variation

The fate model produces a mixture of discrete states. To represent variation within each state, we train a separate conditional flow in the fixed 50-dimensional PCA space. For a training descendant with latent representation *z*_1_ ∈ ℝ^50^ and observed state annotation *S*, draw a Gaussian base point *z*_0_ ~ *N* (0, *I*) and form a point on the linear interpolation *z*_*u*_ = (1− *u*)*z*_0_ + *uz*_1_ for *u* ∈ [0, 1]. The velocity field *v*_*θ*_ (*z*_*u*_, *u, S, h*_*c*_, *t*) is trained by conditional flow matching [19, 37], in the same broad deterministic-transport family as rectified flows [38]:

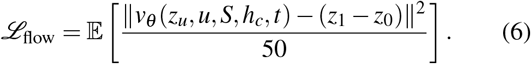

The state condition *S* during training is the descendant’s *observed* annotation. At prediction time no future annotation is available. Instead, the method draws *S* ~ Cat(*p*_*c*_(*t*)), initializes a Gaussian base point and integrates the corresponding conditional velocity field. A fourth-order Runge–Kutta method with 32 steps is used for generation. Sampling the fate before the conditional flow implements an explicit mixture over fates; simply averaging state-specific velocity fields with prior mixture weights need not reproduce this distribution [19, 39, 40].

There are two different time variables. The biological horizon *t* appears in the state-mass evolution and conditions the expression generator. The interpolation coordinate *u* controls movement from base noise to a generated expression profile; it is not elapsed biological time. Generated profiles live in the training-fitted PCA space, and inverse PCA is used only for diagnostics in normalized expression coordinates. In particular, this model does not specify a probability distribution over raw, integer UMI counts.

## 3 Results

### 3.1 Prospective fate prediction from early lineage measurements

We first evaluated the method on the in-vitro LARRY hematopoiesis data, in which lineage barcodes connect observations at days 2, 4 and 6 [1]. The task was to predict the distribution of later labelled descendants using only cells measured on day 2. Eleven annotated states were considered: one undifferentiated population and ten mature categories. To avoid information leakage through multi-barcode cells, connected components of the barcode–cell graph, rather than nominal barcode strings, formed the units of splitting. Five fixed partitions placed 70% of components in training, 15% in validation and 15% in testing. These partitions quantify sensitivity to clone sampling within one experiment; they do not represent five biological replicates.

This evaluation deliberately differs from methods that can examine the complete future dataset while inferring a trajectory. We therefore compared LineageFlow with several predictor classes under a matched early-only information boundary: an empirical prior, a direct conditional mixture flow, a capacity-matched time-softmax predictor, continuous transport and the growth-aware population model PRESCIENT [17]. We also constructed training-only inductive adaptations of CoSpar, CellRank 2 and CLADES, denoted CoSpar-I, CellRank 2-I and CLADES-I [4, 6, 8]. Their scores assess these specific adaptations, *not* the published methods under their full native observation protocols. Representations, tuning and neighborhood selection were restricted to training data (Methods).

### 3.2 Growth correction improves long-term fate prediction

LineageFlow gave the lowest reported Brier loss at both evaluated horizons (Table 2). The distinction was clearest on day 6, when Brier loss was 0.330 compared with 0.346 for CLADESI, 0.355 for the time-softmax predictor and 0.362 for the direct conditional mixture. These changes correspond to absolute reductions of 0.016, 0.025 and 0.032, respectively. The day-6 log loss also favored LineageFlow (1.204 versus 1.258 for CLADES-I), suggesting that the benefit was not specific to the squared-error form of the Brier score. The day-4 gaps were smaller, consistent with a mechanism whose influence accumulates over a longer interval.

**Table 1.**
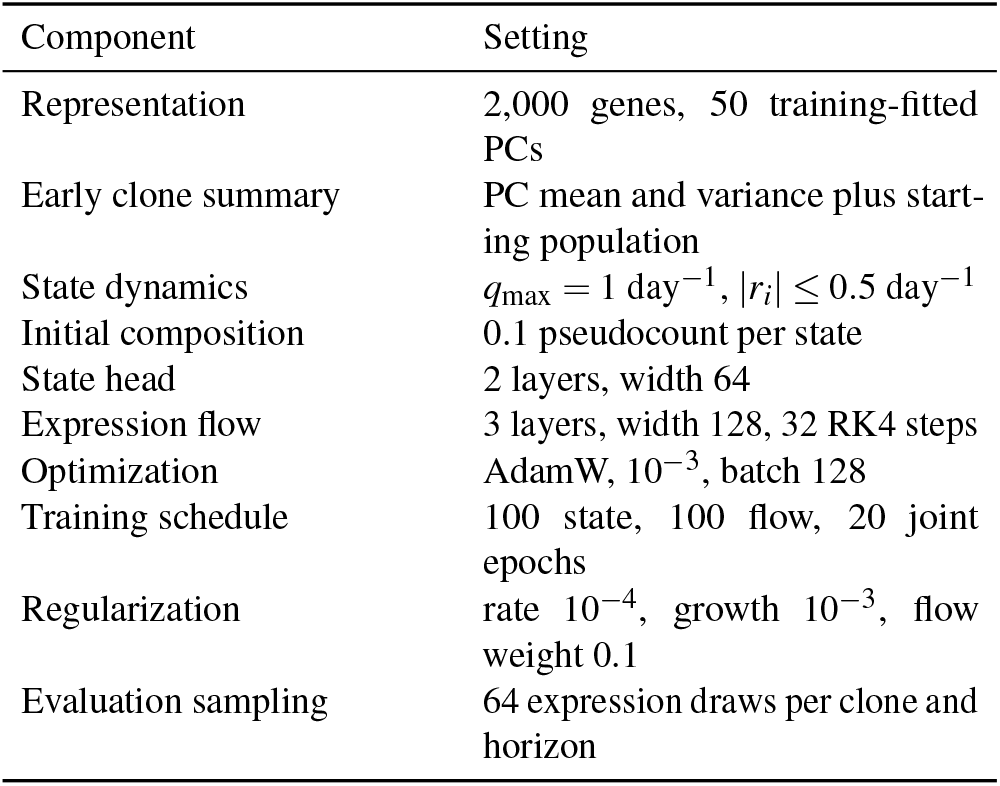
Implementation settings shared across matched neural comparisons. Future test cells do not enter representation fitting, neighborhood construction or tuning.

**Table 2.** Clone-disjoint finite-horizon prediction on the LARRY in-vitro time course. Lower values are better. Day-6 expression scores use 50 training-fitted principal components. I denotes a training-only inductive adapter for a method whose native protocol is not prospective on unseen early clones. Values are means over five clone partitions.

| Method | Brier, day 4 | Brier, day 6 | Log loss, day 6 | Energy, day 6 | ECE, day 6 (%) |
| --- | --- | --- | --- | --- | --- |
| Empirical prior | 0.427 | 0.452 | 1.487 | 0.889 | 9.0 |
| Conditional mixture flow | 0.333 | 0.362 | 1.286 | 0.729 | 5.7 |
| Time-softmax | 0.324 | 0.355 | 1.274 | 0.730 | 5.3 |
| OT-CFM | 0.342 | 0.370 | 1.306 | 0.723 | 6.3 |
| CoSpar-I | 0.327 | 0.354 | 1.299 | 0.804 | 5.8 |
| CellRank 2-I | 0.336 | 0.367 | 1.335 | 0.818 | 6.1 |
| PRESCIENT | 0.322 | 0.353 | 1.291 | 0.744 | 6.0 |
| CLADES-I | 0.316 | 0.346 | 1.258 | 0.781 | 5.5 |
| **LineageFlow** | **0.309** | **0.330** | **1.204** | **0.696** | **4.2** |

To assess whether the ranking was driven by different sets of held-out clones, we compared model scores on the same components. The paired LineageFlow-minus-CLADES-I Brier difference was −0.016, with a 95% clone-component bootstrap interval from −0.024 to −0.008 (Fig. 2b). The corresponding average differences were −0.025 for time-softmax and −0.032 for the conditional mixture. These uncertainty estimates describe resampling of components within the heldout partitions and should not be interpreted as independent experimental or donor-level replication.

The continuous-expression metric provided a complementary comparison. The energy score was 0.696 for LineageFlow, 0.723 for OT-CFM, 0.729 for the conditional mixture flow and 0.781 for CLADES-I. Thus, an improvement in state composition did not come at the expense of the latent-expression score. The state and expression components were subsequently disentangled by matched ablations, rather than assuming that a better joint score identifies which module is responsible.

### 3.3 Predictive performance is robust to descendant-count thresholds

Observed lineage size varies substantially across clones. Of 1,264 components with at least two early cells, 914 had at least ten labelled descendants on day 4 and 742 met that requirement on day 6. At day 6, another 236 components had only one to nine labelled descendants, whereas 286 had none recorded. The last group was treated as censored: a missing labelled descendant does not establish that the lineage died or adopted a specific fate. Consequently, the principal prediction metrics apply to the subset with an observed and sufficiently large descendant sample.

We repeated the day-6 comparison after changing the minimum descendant count. A threshold of five retained 865 components; a threshold of 20 retained 488. Across this range the Brier advantage over CLADES-I was between 0.011 and 0.016 (Fig. 2c). The direction of the result therefore did not depend on the default threshold of ten. This sensitivity check does not, however, recover fates for completely unobserved descendants or remove possible selection bias arising from state-dependent capture and survival.

### 3.4 Rare cell states exhibit reduced prediction and expression accuracy

Mean performance can conceal poor estimates for infrequent outcomes. We therefore separated states representing less than 5% of labelled training descendants from more common states, using training-frequency definitions within each partition. The expected calibration error (ECE), a binned diagnostic for probability calibration [41], was 3.6% for common states and 6.8% for rare states, compared with 4.2% pooled (Fig. 2d). This difference is important because a low overall Brier score does not by itself guarantee reliable probabilities for rare fates.

We also assessed whether generated transcriptional profiles remain consistent with their sampled states. A state classifier fitted only to training descendants obtained a macro-F1 of 0.90 when applied to generated common-state profiles and 0.76 for rare-state profiles. The Spearman correlations between generated and observed state-specific marker means, computed after inverse PCA in normalized expression space, were 0.92 and 0.84, respectively. State-conditional energy loss was 0.674 for common states and 0.743 for rare states (Fig. 2e). These measurements support state-conditioned expression generation at the resolution evaluated while revealing larger errors for rare populations. They do not establish raw-count realism, because the decoder produces PCA-space profiles rather than integer transcript counts.

### 3.5 LineageFlow improves longer-horizon fate prediction in cord-blood data

We next asked whether the relative predictive advantage persisted in a different lineage experiment. Using the cord-blood dataset distributed with CLADES [6], we retrained each model with an early-only observation contract and evaluated subsequent observations at days 10 and 17. LineageFlow achieved day-17 Brier loss of 0.301, compared with 0.316 for CLADESI and 0.329 for time-softmax (Table 3). Its day-17 ECE was 4.7%, compared with 5.4% and 5.8% for those alternatives. Similar ordering at day 10 and day 17 indicates that the ranking was not specific to the LARRY time grid.

**Table 3.** Separately retrained cord-blood evaluation using days 3, 10 and 17. Lower values are better. This analysis tests within-dataset replication on a distinct lineage experiment.

| Method | Brier, d10 | Brier, d17 | ECE d17 (%) |
| --- | --- | --- | --- |
| Conditional mixture | 0.302 | 0.338 | 6.0 |
| Time-softmax | 0.296 | 0.329 | 5.8 |
| CLADES-I | 0.287 | 0.316 | 5.4 |
| **LineageFlow** | **0.276** | **0.301** | **4.7** |

This analysis is a *within-dataset replication after retraining*: it is evidence across distinct lineage experiments, not evidence that a model trained in one experiment transfers without adaptation to a previously unseen donor or culture. That distinction matters when interpreting the breadth of the result.

### 3.6 State dynamics and expression modeling provide complementary benefits

The model contains two sources of flexibility, raising the question of whether its improvement merely reflects extra capacity. A matched two-by-two comparison paired either an endpoint state predictor or the growth-aware CTMC with either a Gaussian expression decoder or the conditional flow (Table 4). With the endpoint predictor fixed, substituting the flow for the Gaussian decoder reduced energy loss from 0.788 to 0.729, whereas Brier changed only from 0.365 to 0.362. Conversely, with the Gaussian decoder fixed, substituting growth-corrected state dynamics reduced Brier from 0.365 to 0.333. The same separation of effects was visible when the conditional flow was retained.

**Table 4.** Matched mechanism controls at day 6. The first four rows form a dynamics-by-decoder factorial.

| Construction | Brier | Energy |
| --- | --- | --- |
| Endpoint head + Gaussian | 0.365 | 0.788 |
| Endpoint head + flow | 0.362 | 0.729 |
| Growth CTMC + Gaussian | 0.333 | 0.757 |
| Growth CTMC + flow | 0.330 | 0.696 |
| Equal-growth CTMC + flow | 0.354 | 0.726 |
| Pooled-rate CTMC + flow | 0.374 | 0.748 |

Two additional restrictions probed the biological interpretation of the state module. Removing differential growth increased day-6 Brier from 0.330 to 0.354. Replacing clone-dependent transition rates with pooled rates increased it to 0.374. These ablations support the usefulness of relative expansion and early-clone context for the stated predictive task; they do not independently establish that the individual fitted transition or expansion rates are uniquely identifiable biological parameters.

### 3.7 Fate predictions are more robust than inferred growth parameters

A constrained transition graph improves interpretability but may not reproduce every biological path. To examine this dependence, we first varied the initial-composition pseudocount, maximum transition rate and allowed range of relative growth. Day-6 Brier remained comparatively stable under moderate changes to these settings (Fig. 3a,b). We then compared the base graph, in which mature states are absorbing with respect to fate transitions, with graphs allowing mature-state switch-ing, a split undifferentiated state or smoothly time-varying rates.

The respective Brier losses were 0.332, 0.328 and 0.329, close to the base model’s 0.330. In contrast, rank correlations between fitted relative-growth parameters and those from the base graph were only 0.62–0.71 (Fig. 3c). The prediction is therefore more robust than a mechanistic interpretation of the inferred rates. In particular, relative growth should be viewed as an effective correction for observed descendant abundance rather than a separate measurement of division or death.

### 3.8 Growth correction improves predictions under unequal expansion

The empirical data do not provide ground-truth division, death and differentiation rates. We therefore examined a controlled five-state system with two differentiation branches,*U* → *MP*→ *M* and *U* → *EP*→ *E* (Fig. 4). The known transition rates were (0.29, 0.21, 0.42, 0.36) day^−1^ and the relative-growth vector was (0.12, 0.19, 0.02, −0.05, −0.28). Clone context biased the two branch rates in opposite directions. The generated descendant observations were sampled from *normalized expected mass*; this is a controlled composition model, not a cell-by-cell stochastic birth–death simulation.

**Figure 4.**
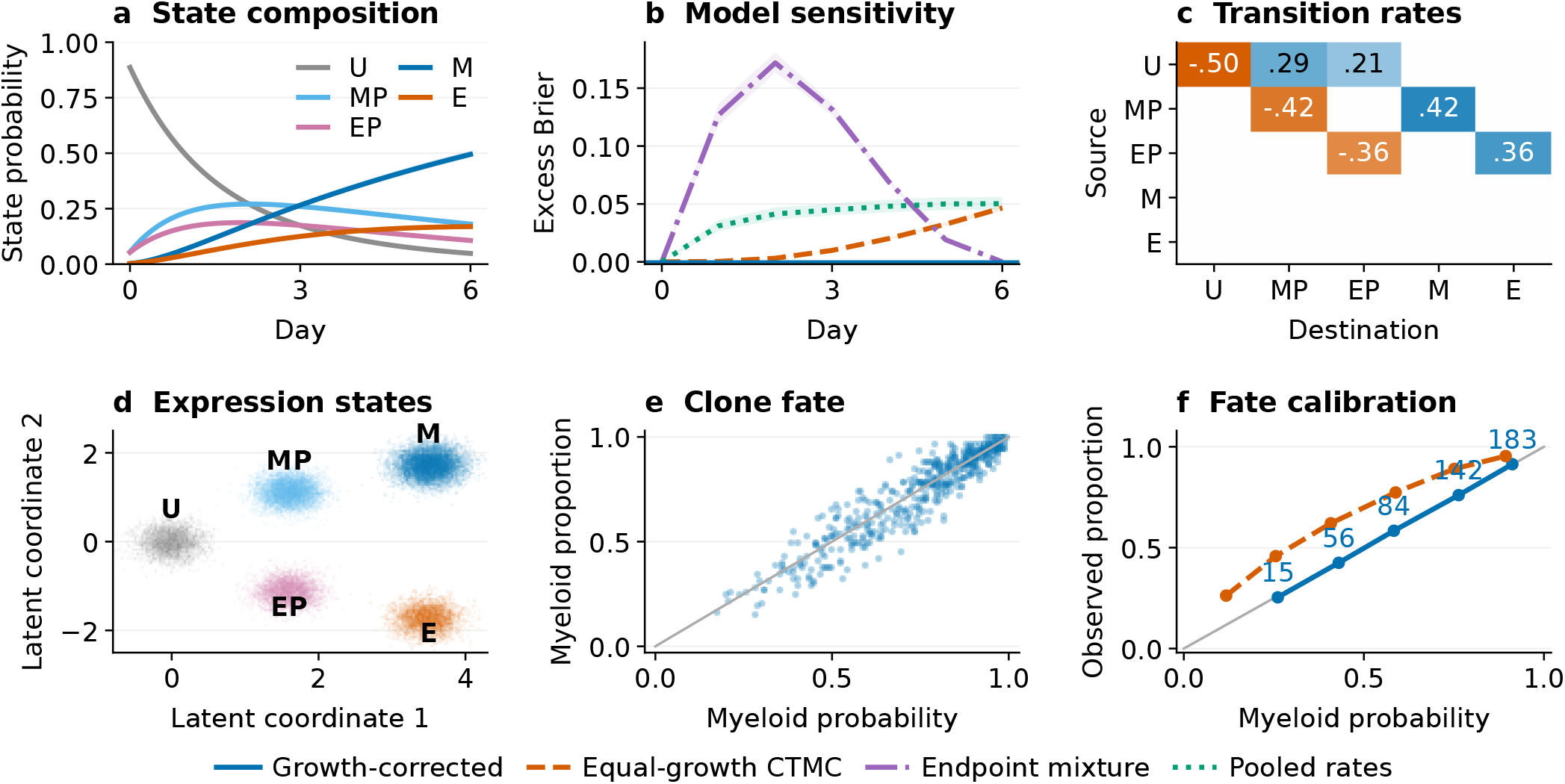
Known-parameter tests separate differentiation and unequal expansion. **a**, Evolution of descendant-state composition in a five-state, two-branch system. **b**, Excess Brier loss relative to the known growth-corrected predictor for alternative model restrictions. **c**, The reference transition generator. **d**, State-conditioned synthetic expression coordinates. **e**, Predicted myeloid fraction for individual clones. **f**, Fate calibration for growth-corrected and equal-growth models. Dense positions along the numerical time grid are not independent observations. Descendants are sampled from normalized expected mass.

Under known parameters, a standard CTMC yields different descendant fractions from the expected-mass model when the terminal states expand at unequal rates. The discrepancy accumulates with time, while setting all state growth rates equal eliminates the difference. An endpoint mixture can match selected observed horizons while deviating between them, and pooled transition rates fail to reflect the clone-specific branch preferences. These experiments explain why growth correction and context dependence are targeted inductive restrictions rather than universally beneficial increases in model complexity.

We also trained reduced versions of the models on simulated clone datasets. For each of five process seeds there were 67 training clones, 14 validation clones and 15 held-out clones. Models observed training horizons 2 and 4, then predicted horizons 1, 3 and 5. The paired mean Brier advantage of the growth-corrected model over its equal-growth counterpart was 0.00283 (sample SD 0.00030 across seeds), but only 0.00001 (sample SD 0.00005) when the data-generating growth rates were equal (Fig. 5c). An omitted mature-state transition introduced after day 3 produced a fitted growth multiplier of 1.39830, despite a true multiplier of one. Thus, even with controlled ground truth, transition-graph misspecification can be absorbed into the effective growth term. These experiments support the prediction mechanism while setting a clear limit on its mechanistic interpretation.

**Figure 5.**
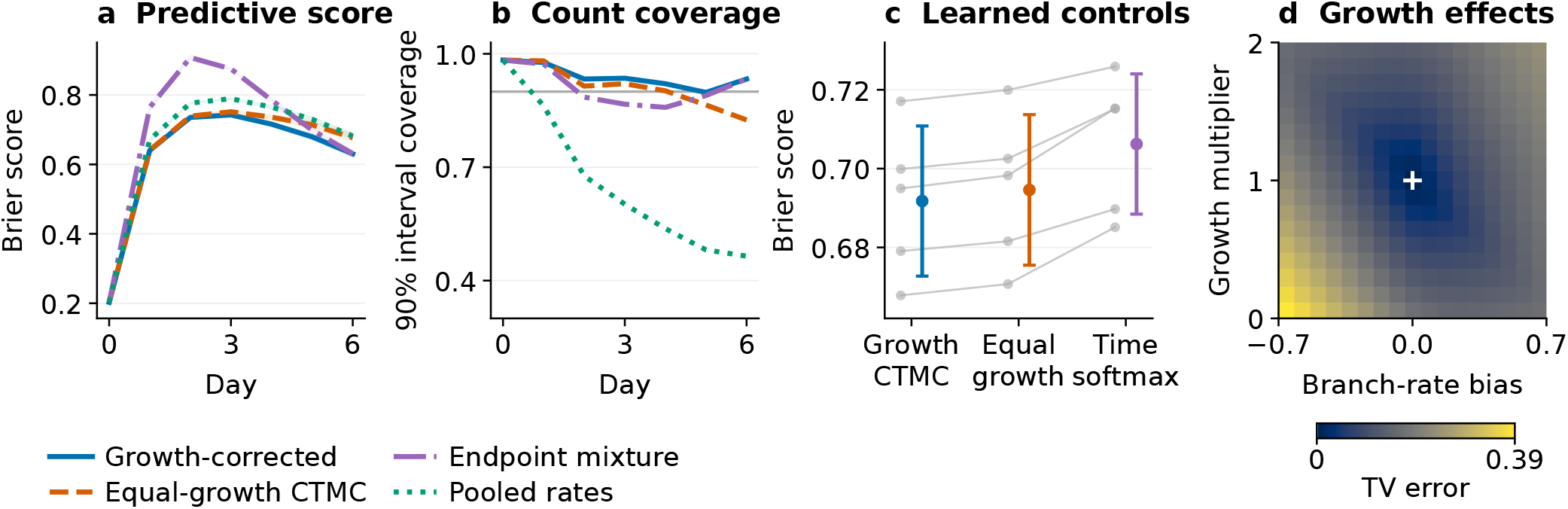
Multi-horizon behavior in the controlled system. **a**, Predictive Brier loss under known parameters. **b**, Coverage of conditional binomial count intervals. **c**, Separately fitted reduced models evaluated on 15 held-out clones at horizons 1, 3 and 5 for each of five process seeds. Lines connect results from the same seed; whiskers denote marginal standard deviations over seeds, not confidence intervals. The mean paired equal-growth-minus-growth Brier difference is 0.00283 (sample SD 0.00030). **d**, Total-variation error under biased branch rates and altered growth multipliers. Panels a, b and d use known parameters; panel c uses learned predictors.

## 4 Discussion

Lineage tracing provides evidence of common ancestry but does not directly observe the trajectories connecting early and late cells. Our results suggest that forecasting the composition of their sampled descendants benefits from distinguishing state transitions from relative expansion, a distinction previously emphasized in population-dynamics inference [16]. This distinction is strongest at the later measured horizons, where differences in descendant accumulation have more opportunity to alter clone composition. In both analyzed lineage datasets, the growth-aware predictor improved proper fate scores relative to matched early-only alternatives, while state-conditioned expression generation provided a complementary gain in transcriptional-distribution scores.

The model’s intended object is a *finite-horizon predictive distribution*, not a reconstructed lineage tree or a uniquely inferred set of physiological rates. Equation 1 propagates expected lineage mass and normalizes it to descendant proportions. It therefore captures one aspect of population dynamics that ordinary probability-conserving CTMC occupancy omits, but it is not a generative model of individual cell divisions, apoptosis, sampling or lineage extinction. Importantly, our five-state simulations share the expected-mass assumptions of the predictor. They help isolate the algebraic role of unequal growth but should not be treated as validation against a realistic stochastic birth–death process, which would require explicit event-level simulation [42]. Such validation, ideally together with explicit capture and survival models, remains a priority.

The approach also has a specific relationship to earlier computational methods. CoSpar, CellRank 2, lineage-informed transport, PRESCIENT and CLADES address complementary inference problems and contribute the methodological context for this work [4–6, 8, 17]. DestinyNet also analyzes lineage-associated fate prediction, but was not part of the matched benchmark here [7]. Some baselines were adapted to the early-clone information boundary, so the resulting matched scores should not be read as comprehensive head-to-head tests of those tools in their native settings. The coupling between the discrete CTMC module and a conditional continuous generator is inherited from Sketch2Colab [20]; the specific extension introduced here is clone-conditioned, growth-corrected descendant prediction with a state-conditioned transcriptional decoder. Distinguishing these contributions is important when interpreting methodological novelty.

Several limits temper the biological interpretation. First, division, death, fate switching, survival and capture can lead to similar changes in normalized descendant fractions. The observation that graph relaxations retain similar Brier losses while changing growth rankings reinforces the non-identifiability of individual kinetic mechanisms. Second, the scored cohort excludes clones with no labelled later descendant and, in the main analysis, those with fewer than ten. Results are therefore conditional on an observable future sample rather than representative of every early lineage. Third, rare states remain less well calibrated than common states, and generated expression is modeled in a 50-dimensional representation of normalized data rather than as an integer UMI-count distribution. Count-based generative modeling and normalization methods highlight technical and biological variability that the present PCA decoder does not model explicitly [43, 44].

The biological evidence is broad enough to test two lineage datasets and several prediction horizons, but it does not establish unrestricted generalization. The five LARRY partitions are not independent biological replicates. The cord-blood comparison retrains on the second experiment rather than transferring a frozen model to an unseen donor. The next important tests are therefore prospective validation under stronger experimental shifts, stochastic branching simulations with imperfect capture, and comparisons in which the same fixed predictive information is available to all candidate methods. Better treatment of small and censored clones would also extend the range of lineages for which calibrated forecasts can be provided.

Taken together, the results support a restricted but useful conclusion: for clone-level fate prediction, it can matter not only *which* states descendants may occupy, but also *how much* those states contribute to the sampled population. A model that represents both effects can offer better finite-horizon fate probabilities without requiring the fitted rates to be interpreted as uniquely measured differentiation or proliferation kinetics.

## 5 Methods

### 5.1 Objective, optimization and prediction protocol

The state module is trained with the clone-weighted negative log probability of observed descendant fractions,

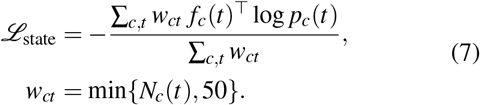

The descendant-count cap prevents a few very large observed clones from dominating the loss. Rate and growth penalties discourage extreme parameter values. The fitting schedule first optimizes the state component, then the expression flow, and finally the shared representation jointly. The model uses AdamW at a learning rate of 10^−3^, with a batch size of 128; the respective stages comprise 100, 100 and 20 epochs. The state head has two layers of width 64, and the expression flow has three layers of width 128. Table 1 records the remaining settings and regularization weights.

For an unseen clone, all early profiles and state assignments are encoded once to obtain *h*_*c*_ and 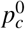. The generator *Q*_*c*_ and growth vector *r*_*c*_ are fixed from that early context and propagated to each requested horizon. Future cells are *not* used to re-estimate a test clone’s context, transition rates, neighborhood or state distribution. A fate is sampled from the predicted mixture and expression is generated conditional on that state and horizon. This is a supervised finite-horizon predictive construction; the training criterion is not the likelihood of an explicitly simulated stochastic branching tree.

### 5.2 LARRY cohort formation and leakage control

The primary evaluation uses the publicly described in-vitro LARRY hematopoiesis time course [1]. Mouse progenitor samples from LSK and LK starting populations were observed on experimental days 2, 4 and 6. Released metadata provide transcriptional profiles, clone barcodes, starting-population labels, sample times, library or replating-well information and mature-state annotations. Replating wells represent branches of the same early culture, not independent donors.

Some cells carry more than one barcode and thereby connect nominal barcode groups. We construct a bipartite cell– barcode graph and use its connected components as indivisible splitting units. Components must have at least two early sampled cells to enter the early-eligible cohort. The principal horizon-specific score also requires at least ten annotated future descendants: 914 of 1,264 early-eligible components on day 4 and 742 on day 6. On day 6, 236 additional components have one to nine annotated descendants and 286 have no labelled descendant. We do not treat this last category as a zero-probability fate or infer cell death from it.

Five fixed component-level partitions contain 70% training, 15% validation and 15% test components. Repeating the analysis across these partitions measures sensitivity to sampling the available clone components within the same biological experiment. All fitted preprocessing, gene sets, representations, graph or neighbor constructions, conditional comparators and calibrations use training data only. Clones held out for testing remain absent from those steps. The data split is intentionally stricter than a random cell split because related descendants of a clone can otherwise appear on both sides of the evaluation boundary. Training-only feature selection and dimension reduction also follow the general principle of isolating fitted single-cell preprocessing from held-out measurements [45].

### 5.3 Matched baselines and their observation contracts

Baselines were selected to probe different explanations for the result rather than to imply identical native objectives. An empirical-prior model predicts training-set fate frequencies. The conditional mixture flow shares the early encoder and expression flow but predicts categorical fate directly from early context and horizon. Time-softmax uses the same early representation and decoder together with polynomial functions of biological time; it is a direct capacity-matched temporal challenger. OT-CFM tests continuous transport [37]. PRESCIENT is initialized from the early clone and fitted on training-time distributions with its growth correction [17].

CoSpar-I and CellRank 2-I form their maps using training observations only, then map an unseen early clone to predictions averaged over its 15 nearest training contexts [4, 8]. CLADES-I similarly constructs training-only groups of early contexts and transfers the nearest group’s fitted rate model to the held-out clone [6]. The suffix “I” means an *inductive adapter*; these are prospective reinterpretations of methods that otherwise often use broader datasets or different observation settings. Therefore, comparisons are specifically about these adapters under a shared information boundary, not about the published tools’ original transductive performance. Fate-only competitors use a common conditional Gaussian PCA decoder where expression scoring requires one. Neural methods use the same early inputs, training-only target representation and validation budget. Values in Table 2 arise from these matched experiments; none is copied from a published benchmark table.

### 5.4 Probability scores, resampling and cohort sensitivity

For a clone’s categorical prediction *p* and observed descendant proportions *f*, we calculate the multiclass Brier loss [46]

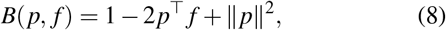

which equals the average of individual-descendant Brier scores within the clone. The corresponding categorical log loss is − *f* ^⊤^ log *p*. Both are proper scoring rules for the observed descendant-state distribution [47]. Scores are aggregated so that each scored clone receives equal weight, rather than weighting the final metric by its number of captured descendants.

Expression-distribution quality is assessed in the fixed training-fitted PCA coordinates using the normalized energy score. For an observed descendant profile *z* and independent generated profiles *Z, Z*^′^, the score is

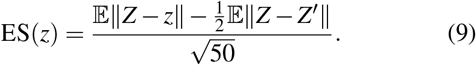

We use 64 generated expression draws per clone and horizon; pairwise estimates exclude matching a sample with itself. The score is averaged within clone before aggregation across clones. A debiased Sinkhorn divergence with fixed cost and regularization is included as a secondary distributional diagnostic [48]. These metrics quantify agreement of latent expression distributions; they are not tests of a count-generating observation model.

For classwise calibration, we use ten fixed bins over clone– state predicted probabilities and compute expected calibration error, a standard but bin-sensitive summary of predictive calibration [41]. States with fewer than 5% of labelled training descendants in a partition are defined as rare, and common and rare summaries are reported separately. Paired uncertainty intervals for score differences resample held-out clone components within partitions while preserving each method’s prediction for the same component. The resulting intervals describe the variation induced by sampled clone components, not variation among independently collected biological donors. Finally, we repeat the day-6 analysis using minimum descendant-future thresholds between 5 and 20, leaving components with no observed future cells censored. This checks threshold sensitivity but cannot correct unknown capture probabilities.

### 5.5 Expression consistency and structural sensitivity

We evaluate generated profiles in three complementary ways. First, a reference cell-state classifier is trained using only observed training descendants and checked against held-out real cells before examining generated profiles. Its predicted state labels on synthetic samples provide a measure of agreement with the sampled conditioning state. Second, marker-gene sets that discriminate states are selected from training cells. Generated PCA profiles are mapped back using the training inverse PCA transform, and their normalized marker-expression means are compared with observed state-specific means by Spearman correlation. Third, state-conditional energy scores quantify distributional agreement separately for common and rare categories. These diagnostics assess different properties and should not be reduced to a single claim of biologically faithful transcriptome generation.

We examine whether the assumed state graph changes forecast quality or the interpretation of fitted parameters. Besides the base graph with ten absorbing mature-state categories, we consider a regularized version permitting mature-state switching, a representation that splits the undifferentiated category, and smoothly time-varying state rates. We compare day-6 Brier scores and rank correlations of estimated relative-growth parameters. Separate sweeps vary the initial-state pseudocount, maximum transition-rate bound and allowed growth range while holding partitions and the remainder of the training procedure fixed. Defaults remain fixed even if a tested alternative produces a slightly lower held-out score.

### 5.6 Independent lineage dataset and controlled branching experiments

The secondary experiment uses the cord-blood lineage-tracing dataset described and distributed with CLADES [6]. It contains observations on days 3, 10 and 17. The same prospective early-only comparison is applied after *separate retraining* on this dataset. The experiment assesses replication of relative model ranking under a different lineage system and time grid; it is not a test of frozen-model transfer to unseen donors.

To isolate state transitions and expansion under known parameters, we also use the five-state branching topology *U* → *MP* → *M* and *U* → *EP* →*E*. Rates along these edges are (0.29, 0.21, 0.42, 0.36) day^−1^ and the reference relative-growth terms are (0.12, 0.19, 0.02, −0.05, −0.28). Clone context shifts the two lineage-branch rates in opposing directions. Synthetic descendants at each observation horizon are sampled from the normalized expected mass given by Eq. 1. Consequently, unlike an event-level stochastic simulation of a birth–death process [42], this controlled experiment does *not* simulate individual stochastic birth, death and observation events; coverage in Fig. 5b is conditional on its specified sampling model. The known-parameter comparisons isolate differences among equal-growth CTMC occupancy, context-pooled rates, endpoint mixtures and growth-corrected expected-mass propagation.

In a separate reduced-model experiment, each of five process seeds gives 67 training clones, 14 validation clones and 15 held-out clones. The training horizons are days 2 and 4, and the evaluated horizons are days 1, 3 and 5. We compare paired clone-level scores for growth-corrected and equal-growth variants within each seed, and report means and sample standard deviations across those five seeds. An additional structural-error experiment introduces an unmodeled mature-state transition of 0.15 day^−1^ after day 3. This identifies how an incorrect transition graph can distort the inferred growth multiplier even when the composition forecast remains plausible.

